# FGTpartitioner: A rapid method for parsimonious delimitation of ancestry breakpoints in large genome-wide SNP datasets

**DOI:** 10.1101/644088

**Authors:** Tyler K. Chafin

## Abstract

1. Partitioning large (e.g. chromosomal) alignments into ancestry blocks is a common step in phylogenomic analyses. However, current solutions require complicated analytical assumptions, or are difficult to implement due to excessive runtimes and unintuitive documentation. Additionally, most methods require haplotype phasing, which is often intractable for non-model studies.
2. Here, I present an efficient and rapid solution for partitioning large genome alignments into ancestry blocks, which better accommodates non-model diploid organisms in that phasing information is not required *a priori*.
3. FGTpartitioner processes a full-chromosome alignment orders of magnitude faster than alternative solutions, while recovering identical results, both via algorithmic improvements and the use of native parallelization.
4. FGTpartitioner provides a means for simple and rapid block delimitation in genome-wide datasets as a pretext for phylogenomic analysis. It thus widens the potential uses for researchers studying phylogenetic processes across large, non-model genomes. Complete code and documentation for FGTpartitioner are available as an open-source repository on GitHub: https://github.com/tkchafin/FGTpartitioner

## Introduction

Inferring genome-wide variation in localized ancestry is a critical step in reconstructing species trees (Springer & Gatesy, 2018), and as such has become a major goal in phylogenomic studies utilizing whole-genome datasets. Analyzing this variation is necessary to understand processes such as horizontal transfer and hybridization, and has led to insights into adaptive introgression (Fontaine et al., 2015), genome-wide selection (Sabeti et al., 2002), disease susceptibility (Seldin, Pasaniuc, & Price, 2011), and ancestral human demographics (Sankararaman et al., 2014). However, partitioning such alignments is a non-trivial task, particularly for researchers utilizing non-model organisms for which limited genomic reference data exists.

Multiple approaches have been proposed for delimiting ancestry blocks in genomes (i.e. establishing recombination breakpoints), which generally fall into one of two categories: those which require dense or phased genotypic data (Liu et al., 2013); and those with complex analytical assumptions which require the definition of informative prior probability distributions and are computationally intensive (Dutheil et al., 2009). Both conditions are problematic for genome-scale studies of non-model diploid organisms, where large-scale resequencing and phased reference data are unavailable, and genomes are often sequenced at low coverage.

I here describe a solution, FGTpartitioner, which is specifically designed for use with non-model genomic data without the need for high-quality phased reference data or dense population-scale sampling. FGTpartitioner delimits chromosome scale alignments using a fast interval-tree approach which detects pairwise variants which violate the four-gametes assumption (Hudson & Kaplan, 1985), and rapidly resolves a most parsimonious set of recombination events to yield non-overlapping intervals which are both unambiguously defined and consistent regardless of processing order. These sub-alignments are then suitable for separate phylogenetic analysis, or as a ‘first pass’ which may facilitate parallel application of finer-resolution (yet more computationally intensive) methods.

## Program Description

For ease of application, inputs are required to follow the widely used VCF format (Danecek et al., 2011). Users may provide parameter settings as arguments in the command-line interface which can restrict block delimitation to a certain chromosome (<-c> flag), with the option to additionally target a region via start (<-s>) and end (<-e>) coordinates. Parallel computation is also possible (<-t>) for particularly large alignments. After parsing user-inputs, the workflow of FGTpartitioner is as follows:

1. For each SNP, perform four-gamete tests sequentially for rightward neighboring records, up to a maximal physical distance (if defined; <-d>) and stopping when a conflict (=’interval’) is found. Intervals are stored in a self-balancing tree. When using multiprocessing (<-t>), daughter processes are each provided an offset which guarantees a unique pairwise SNP comparison for each iteration
2. Merge interval trees of daughter processes (if <-t 2 or greater>)
3. Assign rank *k* per-interval, defined as the number of SNP records (indexed by position) spanned by each interval
4. Order intervals by *k*; starting at min(*k*), resolve conflicts as follows: For each candidate recombination site (defined as the mid-point between SNPs), compute the depth *d* of spanning intervals. The most parsimonious breakpoint is that which maximizes *d*

These algorithm choices have several implications: indexing SNPs by physical position guarantees that the same recombination sites will be chosen given any arbitrary ordering of SNPs; and defining breakpoints as physical centerpoints between nodes means that monomorphic sites will be evenly divided on either side of a recombination event. Because monomorphic sites by definition lack phylogenetic information, they cannot be unambiguously assigned to any particular ancestry block, thus my solution is to evenly divide them.

Heterozygous sites in diploid genomes are dealt with in multiple ways. By default, FGTpartitioner will randomly resolve haplotypes. The user can select an alternate resolution strategy (<-r>) which will either treat a SNP pair as failing if any resolution meets the four-gamete condition, or as passing if any possible resolution passes (i.e. the ‘pessimistic’ and ‘optimistic’ strategies of Wang *et al.*, 2010).

## Benchmarking and comparison to other software

Performance benchmarking was conducted FGTpartitioner and similar methods using publicly available data from the NCBI SRA. I ran tests using a 4-taxon alignment of *Canis lupus* chromosome 1 (~120 Mb), generated from raw reads (*C. lupus*: SRR7107787; *C. rufus*: SRR7107783; *C. latrans*: SRR1518489; *Vulpes vulpes*: SRR5328101-115). Reads were first aligned with bowtie2 (Langmead & Salzberg, 2012) and subsequently genotyped using the HaplotypeCaller and best practices from GATK (McKenna et al., 2010). Tests were all conducted within a 64-bit Linux environment, on machines equipped with dual 8-core Intel Xeon E5-4627 3.30GHz processors and 265GB RAM.

Memory usage for processing a full alignment with FGTpartitioner on 16 cores peaked at 18GB (with less memory required when using fewer cores). Total runtime scales linearly with the maximum allowable physical distance between SNPs to check for four-gamete conditions, with <-d 250000> taking ~50 minutes and <-d 100000> taking 36 minutes to process an alignment comprising >2 million variants. Increasing number of cored offer diminishing returns, with 1, 2, 4, 8, and 16 cores taking 39, 20, 12, 10, and 9 minutes, respectively (to process a 1 million base subset of the 4-taxon chromosome 1 alignment). Runtimes also scale linearly with dataset size (in total variants). Haplotype resolution strategy (random, pessimistic, or optimistic) was not found to impact runtimes.

BA3-SNPS-autotune was used to find optimal mixing parameters for each run, with exploratory analyses employing 10,000 MCMC generations in length. A maximum of 10 exploratory analyses were conducted for each data file. The number of repetitions required to find optimal mixing parameters was recorded for each, and mixing parameters verified to produce adequate MCMC acceptance rates (i.e., 0.2 < acceptance rate < 0.6). All tests were performed on a computer equipped with dual Intel Xeon E5-4627 3.30GHz processors, 265GB RAM, and with a 64-bit Linux environment. Since neither program is multithreaded, a single processor core was used per analysis.

For comparison, I attempted to delimit recombinatorial genes using an alternate method based on the four-gamete test, RminCutter.pl (https://github.com/RILAB/rmin_cut). Because RminCutter does not handle diploid data, I created pseudohaplotypes by randomly resolving heterozygous sites (pseudoHaploidize.py; https://github.com/tkchafin/scripts). I also ran the MDL approach (Ané, 2011), which uses a parsimony criterion to assign breakpoints separating phylogenetically homogenous loci, while penalizing for the number of breakpoints. Neither MDL nor RminCutter could complete an analysis on the full chromosome 1 dataset in a reasonable time span (capped at 10 days), so I first divided the alignment into blocks of ten-thousand parsimony-informative sites each (as these determine runtime, not total length). This produced 21 sub-alignments. MDL took 24 hours across 48 cores to complete a single 5 Mb segment, thus was not further explored. RminCutter.pl took an average of 22 hours per segment (single-threaded). Of note, RminCutter and FGTpartitioner yielded identical results when using comparable settings restricted to a pseudo-haploid dataset. FGTpartitioner thus offers an efficient and methodologically simple solution for ancestry delimitation in non-model diploid organisms with large genomes.

## Conclusion

FGTpartitioner has several advantages over similar methods: 1) algorithmic and performance enhancements allow it to perform orders of magnitude faster, thus extending application to larger genomes; and 2) the flexibility of diploid resolution strategies precludes the need for haplotype phasing *a priori*. Validation using empirical data indicated the suitability of FGTpartitioner for highly distributed work on high-performance computing clusters, with parallelization easily facilitated by built-in options in the command-line interface. Additionally, runtime and memory profiling indicate its applicability on modern desktop workstations as well, when applied to moderately sized datasets. Thus, it provides an efficient and under-friendly solution to alignment pre-processing for phylogenomic studies, or as a method of breaking up large alignments in order to efficiently distribute computation for more rigorous recombination tests.

## Acknowledgements

I would like to thank vonHoldt *et al.* (2016) and Kukekova *et al.* (2018) for making their raw datasets available via the NCBI SRA, and the Arkansas High Performance Computing Center and my Ph. D. advisors Drs. Michael and Marlis Douglas for access to computational resources which I used for developing and benchmarking this program.

